# The Whole Proteome, Phosphoproteome, and Glycoproteome Landscape of Pan-Cancer Cell Lines Profiled by Mass Spectrometry and Reverse Phase Protein Array

**DOI:** 10.1101/2024.10.29.620541

**Authors:** Shi Wenhao, He Tianlong, Nan Wang, Annan Qian, Yuqiao Liu, Tang Shaojun, Zhu Yiying

## Abstract

Mammalian cancer cell lines are essential model systems in biomedical research. We conducted multi-level proteomics analyses on 54 widely used cancer cell lines derived from various tissue-of-origins using two prominent proteomics technologies: mass spectrometry (MS) and reverse-phase protein array (RPPA). Our analysis identified 10,088 proteins, 33,609 phosphorylation sites across 7,289 phosphoproteins, and 56,350 site-specific glycans on 16,296 glycosylation sites from 5,966 glycoproteins, along with 305 drug-relevant protein and phosphoprotein targets. Our results reveal both consistent and distinct patterns in protein expression and modification between MS and RPPA, underscoring their complementary strengths as discovery tools. Additionally, we identified protein features that distinguish tissue origins across different cell line lineages. This dataset supports model system selection for drug target-related studies in vitro and provides valuable insights into key signaling pathways. Overall, this comprehensive resource enables new opportunities for exploration in cancer biology and offers significant value to research communities focused on biomarker profiling, drug target discovery, and understanding mechanisms across diverse cancer types.

## Introduction

Human cancer cell lines serve as essential models for uncovering the molecular mechanisms and cellular behaviors associated with oncogenesis and disease progression. Past studies have leveraged various proteomics approaches to establish proteoform-oriented expression profiles in these cell lines, providing critical insights into cancer systems biology. Most of these studies utilized large-scale analyses covering proteome-wide and post-translational modification (PTM) levels, leading to the creation of databases such as the Cancer Cell Line Encyclopedia (CCLE) and the National Cancer Institute’s 60 (NCI-60) cancer cell line collection. Using 10-plex TMT quantitative proteomics, Gygi’s team profiled 375 cell lines across 22 lineages, quantifying over 12,000 proteins across wide dynamic ranges^1^. In line with that, Aebersold’s team conducted quantitative mass spectrometry (MS) proteomics experiments on NCI-60 cancer cells using a SWATH/DIA-based approach, generating over 8,000 unique proteins within which about 3,171 proteins were compared across cell lines^2^. This proteotypic landscape underscores the value of protein-level data in interpreting cellular phenotypes, inferring protein coregulatory networks, and predicting proteo-transcriptomic-informed drug responses. Complementary studies on NCI-60 cell lines employed label-free MS to analyze proteome and kinome profiles, while others have used MS to characterize the phosphoproteome, illuminating the mechanisms of action (MOA) of cancer drugs^3,4^. The DepMap Project further integrates proteomic data with pharmacological profiles, deepening our understanding of molecular vulnerabilities and therapeutic targets in cancer (https://depmap.org/portal/).

In addition to MS-based proteomics, affinity-based protein quantification methods, such as the Reverse Phase Protein Array (RPPA), allow high-throughput quantification of proteins and modified proteoforms using specific antibodies^5,6^. Cell-based RPPA experiments, including those by our team, are commonly used to investigate biological mechanisms under perturbation^7,8^. This technique has characterized NCI-60 cell lines, generating drug response networks and signaling pathway activity profiles that impact cellular fates^9–11^. A key advantage of RPPA is its ability to quantify low-abundance proteins and PTM. For example, Davies and colleagues used a panel of 222 protein features, including total and phosphorylated proteins, to categorize NCI-60 cells into five clusters, each associated with specific mutations linked to drug response^10^. Other studies identified protein-inferred drug response, patterning the activation/phosphorylation states of 135 proteins and defining six core cancer signaling modules related to therapeutic responses^12^. Hundreds of cancer cell lines with more than 200 protein targets have since been profiled by RPPA, with data repositories such as CCLE, PRIDE (https://www.ebi.ac.uk/pride/), and MD Anderson’s Cell Lines Project (MCLP: https://tcpaportal.org/mclp/#/) offering comprehensive datasets that allow researchers to investigate protein functions, drug targets, and biomarkers across cancer types^13–15^. Recent RPPA studies expanded these resources, adding 447 clinically relevant dual-actionable targets in the cancer field^16^.

MS-based proteomics and RPPA each offer distinct advantages in proteomics research. MS-based proteomics digests proteins into peptides, identifies them by comparing MS/MS spectra with theoretical protein sequences, and enables large-scale characterization of the entire proteome and modified forms, whereas affinity detection-based methods such as RPPA employ antibodies to measure protein levels across hundreds of samples simultaneously^5,17–21^. While several studies have integrated MS and RPPA data, most focus on clinical samples. For example, MS and RPPA have been combined to provide quantitative proteomic landscapes of breast cancer tissue^22^. Proteomics data derived from The Cancer Genome Atlas (TCGA) using RPPA (quantifying 150–200 protein forms) have also been compared with global MS proteomics from the Clinical Proteomics Tumor Analysis Consortium (CPTAC, https://proteomics.cancer.gov/programs/cptac), highlighting the role of protein-driven therapy development across multiple cancer types^23^. Despite these advances, deep characterization of cancer cell lines using multiple proteomics methods remains limited, with only a few studies performing such analyses on a systemic level^7^.

Beyond total protein profiling, PTMs, mainly phosphorylation and glycosylation, are critical cellular function governors in cancer^24–27^. Phosphoproteomics, which characterizes site-specific phosphorylation on proteins, is invaluable for uncovering dysfunctional kinase activities and thus aiding the development of therapeutic drugs such as kinase inhibitors^4,17,28,29^. Databases such as PhosphoSitePlus provide extensive data on phosphorylation and other PTM data for cancer cell lines, supporting targeted research (https://www.phosphosite.org/). Similarly, glycoproteomics, which examines glycosylation modifications, has been extensively studied due to its potential as a diagnostic biomarker for cancer and other diseases^30,31^. Recent advances in MS technology have enabled large-scale glycosylation profiling of cancer cell lines^32,33^, offering mechanistic insights into glycosylation on a global scale during oncogenesis and disease progression.

To address the gap in multi-dimensional proteomic analysis of cancer cell lines, we applied label-free MS together with RPPA to obtain quantitative proteomics data on 54 representative cancer cell lines, predominantly from NCI-60. This study provides a comprehensive proteomics landscape, incorporating the total proteome, phosphoproteome, and glycoproteome, which reinforces our understanding of protein-level regulation in cancer cells. By comparing MS and RPPA datasets, our study underscores the complementary strengths of these two methods: MS offers a broad view of protein abundance and PTMs, while RPPA facilitates targeted quantification of specific proteins and PTMs, even at low abundances. This combined approach highlights the value of multi-dimensional proteomic data in discovering therapeutic targets, identifying biomarkers for cancer subtyping, and predicting cellular responses, supporting advances in precision oncology.

## Methods

### Protein extraction and trypsin digestion for MS analysis

Cells were minced and lysed in lysis buffer (8 M urea, 100 mM Tris hydrochloride, pH 8.0) containing protease and phosphatase inhibitors (Thermo Scientific) followed by 1 min of sonication (3 s on and 3 s off pulse, amplitude 25%). The lysate was centrifuged at 14,000 × g for 10 min, and the supernatant was collected as whole tissue extract. Protein concentration was determined by the bicinchoninic acid (BCA) protein assay. Extracts from each sample (1 mg proteins) were reduced with 10 mM dithiothreitol at 56°C for 30 min and alkylated with 10 mM iodoacetamide at room temperature (RT) in the dark for an additional 30 min. Samples were then digested using the filter-aided proteome preparation (FASP) method with trypsin. Briefly, samples were transferred into a 30 kD Microcon filter (Millipore) and centrifuged at 14,000 × g for 20 min. The precipitate on the filter was washed twice by adding 300 μL washing buffer (8 M urea in 100 mM Tris, pH 8.0) into the filter and centrifuged at 14,000 × g for 20 min. The precipitate was resuspended in 200 μL 100 mM NH4HCO3. Trypsin with a protein-to-enzyme ratio of 50:1 (w/w) was added to the filter. Proteins were digested at 37°C for 16 h. After tryptic digestion, peptides were collected by centrifugation at 14,000 × g for 20 min and dried in a vacuum concentrator (Thermo Scientific). 5% peptides were used to detect the proteome, and 95% peptides were used to prepare phospho-peptides with Fe-NTA enrichment kit (Thermo Scientific, A32992).

### Enrichment of phosphopeptides and glycopeptides

The peptide sample was fully dissolved in 100 µL of 80% acetonitrile and 0.1% TFA solution following vortex mixing. Subsequently, the precipitation was removed by centrifugation at 16,000 × g for 10 min. The enrichment column was retrieved from the kit (Thermo Scientific, A32992), and the preservation solution was removed by centrifugation at 1,000 × g for 1 min. This was followed by the addition of 100 µL of 80% acetonitrile and 0.1% TFA solution, which was gently mixed and incubated at room temperature for 3 min. After incubation, the solution was removed by centrifugation at 1,000 × g for 1 min, and the peptide solution supernatant was added to the column. The mixture was incubated at room temperature for 30 min, with gentle mixing every 10 min. Following the incubation period, the peptide solution was removed by centrifugation at 1,000 × g for 1 min. To wash the column, 100 µL of 80% acetonitrile and 0.1% TFA solution was added, and the washing solution was removed by centrifugation, repeating the wash step three times. A clean 1.5 mL Eppendorf tube was then placed beneath the enrichment column, and 100 µL of 50% acetonitrile and 9% ammonia solution was added for elution. The elution was collected by centrifugation at 1,000 × g for 1 min, resulting in the enriched phosphorylated peptide solution. The phosphorylated peptide solution was vacuum-dried, and the peptide powder was stored at −80°C until analysis.

### RPPA sample processing

The Reverse Phase Protein Array (RPPA) was performed following a standardized workflow to ensure consistency and quality. Protein lysates were initially mixed with a sample dilution buffer (comprising 50% glycerol, 4X SDS buffer, and 6 ml of beta-mercaptoethanol) to achieve a final concentration of 1.5 mg/ml. Normalized samples were then further diluted 2-fold in sample dilution buffer, consisting of lysis buffer, 50% glycerol, and 4X SDS buffer with 6 ml of beta-mercaptoethanol in a 3:4:1 ratio. Five serial dilutions (1, 1/2, 1/4, 1/8, 1/16) were performed using automated liquid handling workstations (Tecan Fluent series). The processed samples were loaded into low-binding 384-well plates (Molecular Devices) and then deposited onto nitrocellulose-coated glass slides (Grace Bio-Labs ONCYTE superNOVA) using a Quanterix 2470 solid pin contact printer.

To maintain quality control (QC), on-slide controls, including treated and untreated cell lines as well as a lysate mixture from various cell lines and tonsil tissue, were applied for staining and quantitative QC checks. Approximately 400 identical slides were prepared for the study.

Each slide was then processed for colorimetric signal quantification using a validated panel of 305 antibodies, targeting 227 total proteins and 78 phosphoproteins or other post-translational modifications (PTMs) (detailed in Supplementary S1). The slides were blocked with Re-Blot (Millipore) at room temperature, followed by I-block (Fisher) and antigen retrieval with hydrogen peroxide (Fisher). Sequential blocking steps with avidin, biotin, and protein block (DAKO) were conducted before a 1-hour primary antibody incubation at room temperature. Secondary antibodies (DAKO) specific for rabbit or mouse were applied, followed by Tyramide Signal Amplification (TSA, Akoya) and DAB colorimetric visualization (DAKO). Staining was fully automated using the DAKO Link 48 Autostainer (Agilent), and the slides were then scanned on a high-throughput slide scanner to capture and analyze colorimetric signals accurately.

### LC-MS/MS analysis

Dried peptide samples were re-dissolved in Solvent A (0.1% formic acid in water) and loaded to a trap column (100 μm × 2 cm, home-made; particle size, 3 μm; pore size, 120Å; SunChrom, USA) with a max pressure of 280 bar using Solvent A, then separated on a home-made 150 μm× 12 cm silica microcolumn (particle size, 1.9 μm; pore size, 120Å; SunChrom, USA) with a gradient of 5-35% mobile phase B (acetonitrile and 0.1% formic acid) at a flow rate of 300 nL/min for 120 min.

The eluted peptides were ionized under 2.2 kV. MS was operated under a data-dependent acquisition (DDA) mode. For detection with Orbitrap Eclipse mass spectrometer, a precursor scan was carried out in the Orbitrap by scanning m/z 300-1,500 with a resolution of 60,000. Then, MS/MS scanning was carried out in the Orbitrap by scanning m/z 200-1,400 with a resolution of 15,000. The most intense ions selected under top-speed mode were isolated in Quadrupole with a 1.6 m/z window and fragmented by higher energy collisional dissociation (HCD) with normalized collision energy of 32%. Max injection time set 40 ms for full scans and 30 ms for MS/MS scans.

Dynamic exclusion time was set as 30 s. For phosphopeptide, a precursor scan was carried out in the Orbitrap by scanning m/z 300-1,500 with a resolution of 60,000. Then, MS/MS scanning was carried out in the Orbitrap by scanning m/z 200-1,400 with a resolution of 30,000. The most intense ions selected under top-speed mode were isolated in Quadrupole with a 1.6 m/z window and fragmented by higher energy collisional dissociation (HCD) with a normalized collision energy of 27%. Max injection time set 30ms for full scans and 54 ms for MS/MS scans.

### MS database search

All the MS data were processed in the Maxquant (V1.6.17.0) platform. Raw files were searched against the human National Center for Biotechnology Information (NCBI) ref-seq protein database (updated on 07-04-2013, 32,015 entries). Mass tolerances were 10ppm for precursor and 0.05 Da for product ions. Up to two missed cleavages were allowed. The data were also searched against a decoy database so that protein identifications were accepted at a false discovery rate (FDR) of 1%. Carbamidomethylation (C) was set in the search engine as a fixed modification; Acetyl (Protein N-term) and Oxidation (M) as variable modifications. For phospho-proteome, the variable modifications included phosphorylation on serine, threonine, and tyrosine.

The qualitative analysis of N-glycopeptides from the MS/MS data was conducted using Byonic software version 5.0.3 (Protein Metrics Inc., USA). Employing full tryptic digestion with allowance for up to two missed cleavages, the precursor mass tolerance was fixed at 10 ppm, whereas the fragment-mass tolerance was set at 0.02 Da. Carbamidomethylation of cysteines was set as a fixed modification, and variable modifications were set as acetyl (N-terminus), methionine oxidation (M), and phospho (S, T, Y). The maximum dynamic modification number was restricted to 2. The analysis utilizes the built-in database 132 N-Glycan of human in Byonic. The false discovery rates (FDRs) at the protein level remained below 1%.

### RPPA analysis

Image data were digitally processed using MicroVigene software (version 5.6.0.8), producing both text (.txt) and image (.tiff) files for each slide. These files were then analyzed using SuperCurve fitting via the R package SuperCurve to produce expression data (rawlog2 files) and quality control (QC) metrics for each slide. Correction factors (CF) were calculated to identify outliers both within and across experiments. For data normalization (to adjust for loading differences), median subtraction was applied: first, each antibody column was median subtracted, followed by median subtraction for each sample row. This resulted in a normalized log2 file, which was squared to create a linear dataset (Normlinear), detailed in Supplementary S2. These processed datasets were then prepared for quantitative comparisons and graphical visualization in downstream analyses.

### Data processing

MS-based proteomics and phosphoproteomics data were analyzed using MaxQuant’s label-free quantitation (LFQ) approach to assess protein and phosphorylation-site abundances. Prior to further processing, proteins, and phosphorylation sites present in less than 5% of samples were filtered to account for the broad lineage diversity of cell lines, thus applying a more lenient threshold than usual. Missing values were imputed with the minimum values from each respective sample. Data were manually examined using the R package “DEP” (v1.16.0). Data visualization and preprocessing were performed with “Seurat” (v5.0.1), using the “NormalizeData” function with the “LogNormalize” method. The R package “COSG” was utilized to identify marker proteins, setting a COSG score threshold of 0.5. For calculating the fold change (FC) of phosphorylation sites, the “FindAllMarkers” function in “Seurat” was applied.

For mass spectrometry-based glycoproteomics, each glycopeptide is annotated with its glycan composition, which includes sialic acid (NeuAc), fucose (Fuc), N-acetylhexosamine (HexNAc), and hexose (Hex). A site-specific glycan is defined as a particular type of glycan located at a specific glycosylation site on a protein. This definition encompasses peptides of varying lengths, including those with missed cleavages and differing charge states of the peptide ions. The abundance of each site-specific glycan is estimated using spectral counting, which involves summing the number of spectra corresponding to all peptide ions that contain this type of glycan at the designated sites.

For RPPA data, antibody probes were mapped to Uniprot Accession IDs of target proteins. In cases of multiple mappings, a representative protein was selected to facilitate comparisons between RPPA and MS data.

To enable comparisons across data modalities, the data were processed systematically for consistency across platforms. For MS proteomics and phosphoproteomics, raw abundance values were log2(x+1)-transformed, followed by averaging replicate values to quantify each cell line. In the case of MS glycoproteomics, site-specific peptide counts for each modification, site, or protein were averaged across replicates for cell line quantification. For RPPA proteomics and phosphoproteomics, mean normalized expression values were calculated from replicates to quantify each cell line. Using these standardized quantified values, correlation analyses were performed to examine relationships across cell lines.

### Functional annotation, Enrichment analysis, and KSEA analysis

GO annotations were performed using the “org.Hs.eg.db” R package (v3.18.0), while KEGG pathway data were accessed through “KEGGREST” (October 11, 2024). GO enrichment analysis was carried out using “clusterProfiler” (v4.10.0).

KSEA (Kinase-Substrate Enrichment Analysis) was conducted using “KSEAapp” (v0.99.0) to estimate kinase activity changes by averaging multiple substrate measurements rather than single-substrate dependence. Fold change (FC) values for phosphorylation sites in each cell line were computed as previously described. Kinase-substrate relationships were sourced from PhosphoSitePlus® (January 15, 2024). A Z-score was calculated by comparing the mean log2(FC) of each kinase’s substrate phosphosites against the mean log2(FC) of all phosphosites in that cell line. Kinases with significant Z-scores (FDR < 0.05) and more than five substrates were considered to exhibit altered activity.

## Results

### Overview of the study

In this study, we investigated 54 widely used tumor cell lines representing various human tissues. Cultured cells were harvested and prepared separately for MS and RPPA proteomics analysis. The MS-based analysis was comprehensive, encompassing the whole proteome, phosphoproteome, and glycoproteome (Fig. 1a). These cell lines were derived from tissues on breast (n=7), esophagus/stomach (n=8), lymphoid (n=8), lung (n=5), ovary/fallopian tube (n=5), large bowel (n=5), and other origins (n=16) (Fig. 1b, Supplemental Table S1).

**Figure 1.**
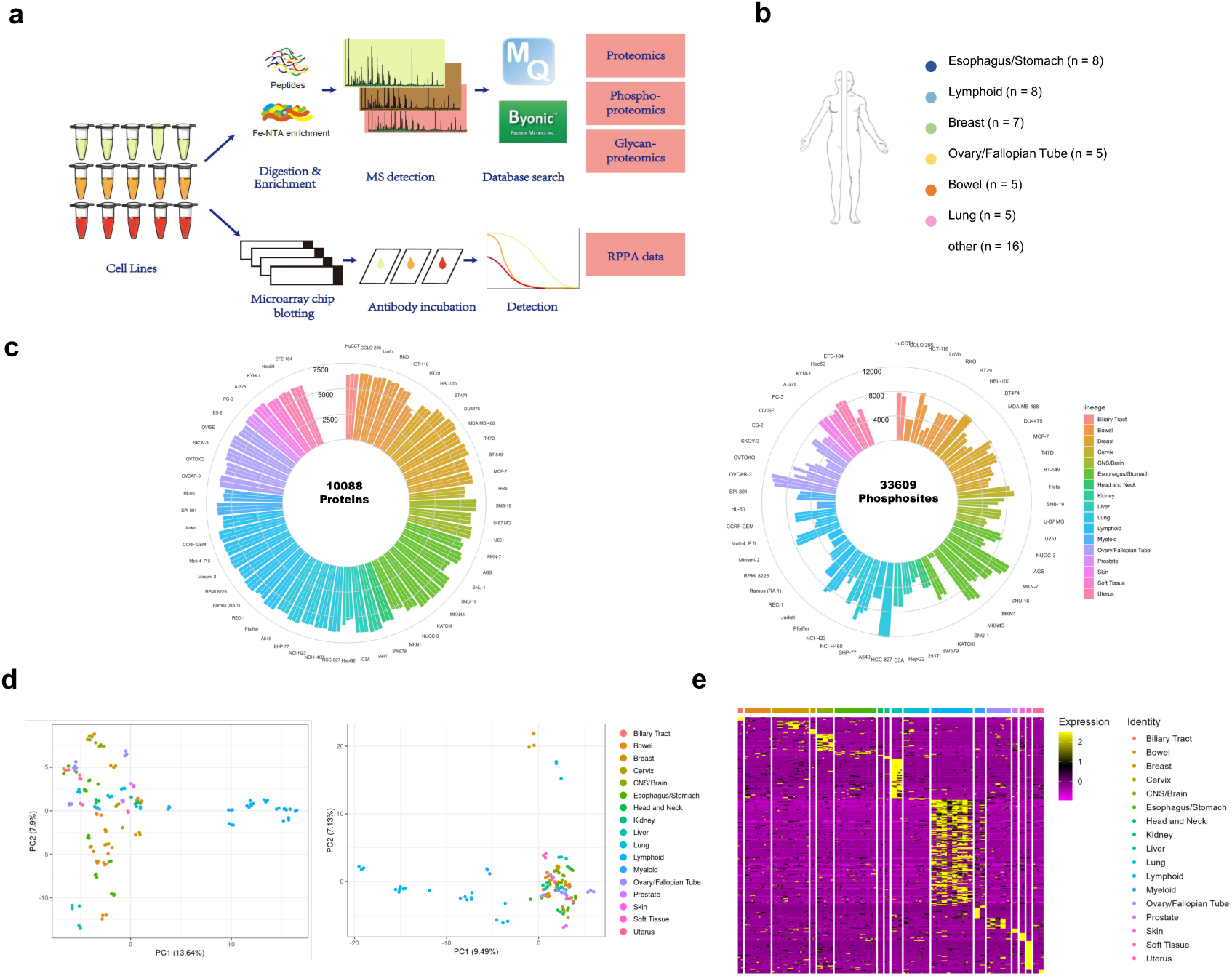
Experimental Design and Overview of the Data (a) Workflow of proteomics analysis of cancer cell lines: Cultivated cells were divided into separate aliquots for MS and RPPA sample preparation. After analysis via mass spectrometry or Array-Pro analyzer, MS data was processed separately for the total proteome, phosphoproteome, and glycoproteome, while RPPA data was processed to measure protein concentrations. (b) Cell line classification: Cell lines are grouped based on their tissue of origin. (c) Number of detections per cell line: The left side shows total proteins detected, and the right side shows phosphosites detected across cell lines. The outer circle lists cell line names, with bars representing the number of detections for each cell line. Bar colors indicate the tissue origin of the cell lines. (d) PCA of cell lines: Principal component analysis (PCA) displays the proteome (left) and phosphoproteome (right) expression profiles for each cell line. Each dot represents a sample, and the percentage value indicates the variance explained. (e) Heatmap of protein expression: The heatmap shows relative protein expression levels across cell lines, with the color scheme representing standardized scores. Yellow indicates over-expressed proteins.

After applying the filtering criteria described in the Methods, we identified 10,088 proteins in the MS-based dataset, with a median of 6,330 proteins detected per sample (Fig. 1c, Supplemental Table S2). In the phosphoproteomics dataset, we detected 9,380 phosphoproteins with 33,609 phosphorylation sites (location probability > 0.75), and median values of ∼3,000 phosphoproteins and 6,000 phospho-sites per sample (Supplemental Table S3). Furthermore, we identified 56,350 site-specific glycans from 16,296 glycosylation sites, spanning 5,966 glycoproteins (Supplemental Table S4). The RPPA panel included 231 whole-protein and 74 phosphosite-specific antibodies (Supplemental Table S5).

Principal component analysis (PCA) of protein expression and phosphorylation (Fig. 1d) demonstrated strong reproducibility in the cell line replicates, confirming the consistency of the data. Phosphorylation reflects the cell’s different states and can vary under external perturbations; in this study, the data represents the basal phosphorylation levels of the cell lines. The analysis revealed significant differences based on tissue origin, with a clear distinction between cell lines derived from nonsolid tumors and solid tumors.

Additionally, a heatmap of whole protein expression (Fig. 1e) highlighted the diversity of expression across cell lines. We identified 292 marker proteins (COSG scores > 0.5) enriched in specific tissues, which demonstrated distinct expression profiles (Supplemental Table S2). These marker proteins were useful in differentiating tissue-specific expression patterns.

This comprehensive proteomics dataset provides a multi-layered understanding of the proteome, phosphoproteome, and glycoproteome across a diverse set of cancer cell lines, offering valuable insights into cancer biology and the molecular underpinnings of tumor-specific behaviors.

### Phosphoproteomics revealed phosphorylation dynamics of cancer cell lines

Phosphorylation, a key post-translational modification (PTM), plays a crucial role in regulating cellular signaling pathways. Oncogenicity is often linked to dysregulated molecular signaling, and understanding phosphorylation patterns can provide insights into cancer mechanisms and therapeutic targets. As of January 2024, the FDA has approved 80 small-molecule kinase inhibitors targeting 24 kinases that regulate protein phosphorylation on serine, threonine, and tyrosine residues, with more therapeutics under development^24,25^. Studying phosphorylation patterns of human cancer cell lines may help estimate the disease mechanisms, select suitable models, and evaluate drug responses^4^. In our MS phospho-proteome analysis, we identified 33609 phosphorylation sites, with 90.9 % of them previously reported in the PhosphoSite Plus database (www.phosphosite.org) (Fig. 2a). Due to the use of immobilized metal affinity chromatography (IMAC) affinity for phosphopeptide enrichment, the distribution of serine, threonine, and tyrosine phosphorylation in this study mirrored the natural occurrence of these PTMs, with serine and threonine being the most common and tyrosine being the least prevalent. In addition, RPPA provided a detailed characterization of specific phosphorylation sites, especially for tyrosine, based on 74 site-specific antibodies, complementing the broader MS-based overview with more targeted data.

**Figure 2.**
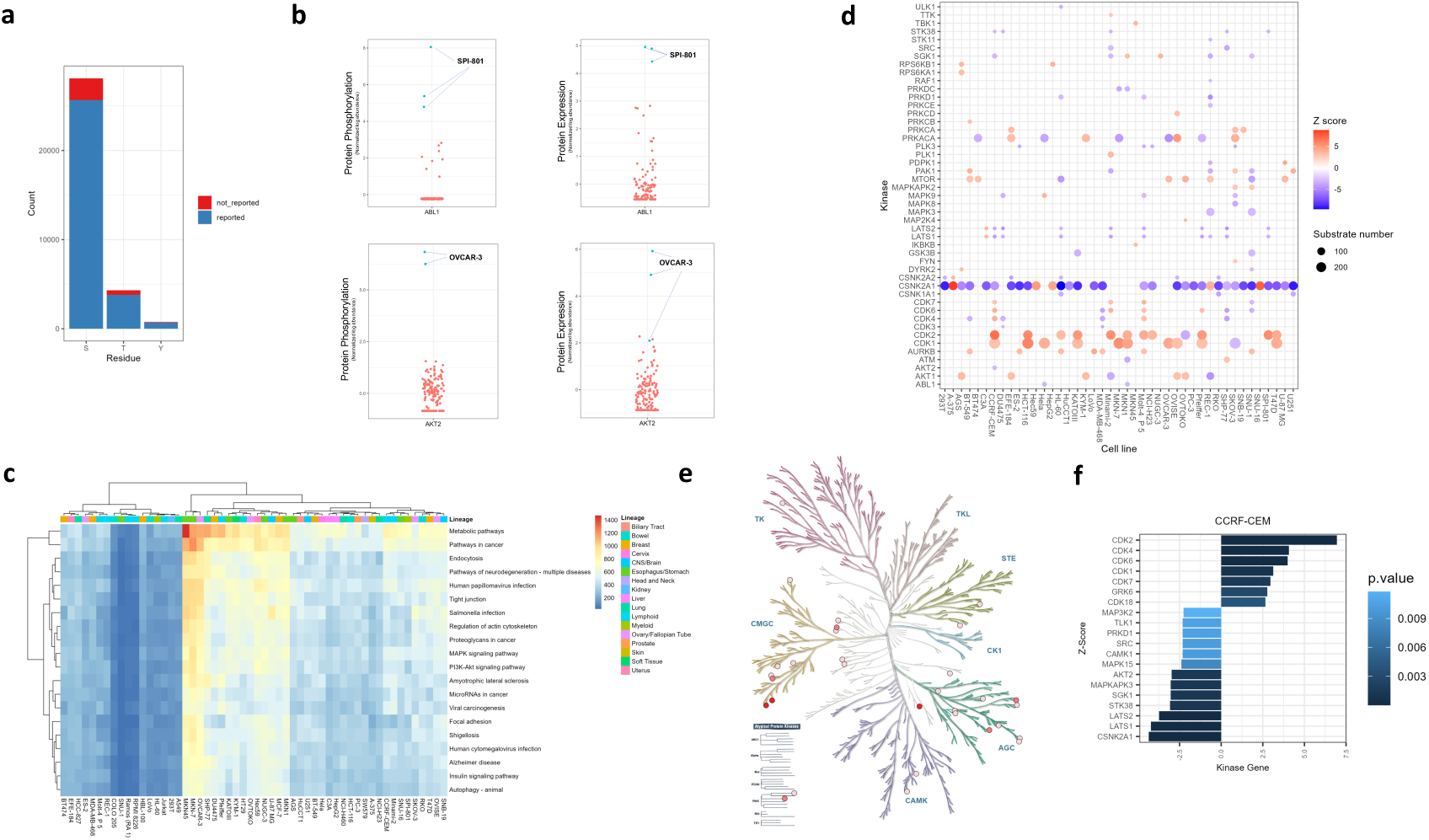
Phosphoproteome Landscape of Cell Lines (a) Distribution of Phosphorylation Types: The modifications (uncurated) are compared against the PhosphoSite Plus database (www.phosphosite.org), with reported modifications shown in blue and unreported modifications in red. (b) Whole Proteome and Total Phosphorylation of ABL1 and AKT2 in Cell Lines: This study presents the abundance levels of both ABL1 and AKT2 proteins, along with their phosphorylation levels in various cell lines. Each dot represents a replicate. Protein abundance is calculated by summing the values of the top three unique peptides, while phosphorylation abundance is determined by the total of all identified phosphopeptides. (c) Heatmap of Signaling Pathways in Cell Lines: The phosphorylation levels of proteins across various cell lines are shown, with the number of detected phosphorylation sites representing these levels. The block colors indicate pathway enrichment, and the top of the heatmap is color-coded to represent the tissue origins of the cell lines. (d) Kinase Activity Inferred from Substrates: Kinase activity is depicted based on the number of phosphorylated substrates across different cell lines. The circle size corresponds to the number of phosphorylated substrates for each kinase, and the color intensity reflects the Z scores, representing the relative abundance of phosphorylated substrates. (e) Kinase Detection in Cell Lines Mapped on the Kinase Tree: This panel maps detected kinases across different categories on a kinase tree. Circle depth represents the number of detections, with darker colors indicating higher detection frequencies. (f) Kinase Activation in CCRF-CEM Cell Line: Kinase activation is illustrated based on the number of substrates phosphorylation and numbers in the CCRF-CEM cell line.

Our data revealed significant variations in phosphorylation levels across different cell lines, which could be useful for identifying abnormal kinase activity. For example, Abelson 1 (ABL1) phosphorylation was notably higher in the chronic myeloid leukemia (CML) cell line SPI-801 (a derivative of K-562) compared to other lines (Fig. 2b). CML is driven by the BCR-ABL1 fusion gene, which results in abnormal tyrosine kinase activity. Targeted therapies, such as imatinib, inhibit BCR-ABL and can induce apoptosis in CML cells^28^. Thus, our data hitlier phosphorylation patterns, which may offer insights for drug development in specific cancer types.

We also generated a heatmap of signaling pathways across different cell lines (Fig. 2c), showing no clear correlation between tissue origin and phosphorylation patterns, suggesting that even cancers from the same tissue may require personalized treatments. Notably, hyperphosphorylation of the PI3K/Akt/mTOR pathway was observed in the gastric cancer cell lines MKN45 and MKN7, consistent with reports that these cell lines are sensitive to inhibitors targeting this pathway from the CancerRxGene database (www.cancerrxgene.org/celllines). Additionally, Akt2 was blotted across cell lines, and its overexpression at the protein level and high phosphorylation level was clearly observed in ovarian cancer cell line OVCAR-3, consistent with the literature picturing its role in cancer cell signaling and potential as a therapeutic target^34–38^ (Fig. 2b).

Further, we performed a kinase-substrate enrichment analysis (KSEA), represented through a dot plot (Fig. 2d) and kinase tree (Fig. 2e). Noteworthy findings included the high activity of CDK4 substrates in the Pfeiffer cell line, a diffuse large B-cell lymphoma line, aligning with reports that CDK4/6 inhibitors are effective against aggressive B-cell lymphomas^39^. In the CCRF-CEM cell line from T-cell acute lymphoblastic leukemia, substrates of CDK4 and CDK6 showed significant phosphorylation (Fig. 2f), reflecting the potential for CDK4/6 inhibitors in this disease^40,41^. We also noted the overexpression of CSNK2A1, a subtype of CK2, across multiple cancer cell lines, including SNU-16 (gastric adenocarcinoma) and HepG2 (hepatoblastoma). CK2 is known to be overexpressed in various cancers, and inhibitors like CX-4945 show therapeutic potential in gastric and liver cancers^42–44^.

In summary, our phosphoproteomics analysis provides a comprehensive view of kinase activities, activated signaling pathways, and the relationships between total protein levels and phosphorylation. These insights are valuable for selecting appropriate cancer cell line models for drug and cellular signaling research, as well as for predicting the sensitivities of cancer cell lines to kinase-targeting therapies. This understanding can help identify key kinase-driven processes and improve the precision of therapeutic interventions in cancer research.

### Glycoproteome landscape of cancer cell lines

Protein glycosylation is a complex post-translational modification (PTM) that significantly influences protein structure, stability, function, and intracellular signaling^45^. Recent advancements in mass spectrometry and computational tools have enabled the large-scale analysis of intact glycopeptides. In this study, we characterized 56350 site-specific glycans at 16296 glycosylation sites from 5966 glycoproteins. The IMAC can enrich glycopeptides carrying sialic acids, as well as phosphorylated glycans such as mannose-6-phosphate and extra acidic amino acid peptides^46^. Among the identified glycans, 49.72% contained sialic acid, 25.11% were high-mannose types, and 25.16% contained fucose (Fig. 3a). This diversity, including glycopeptides found at the same or different protein sites, highlights the complexity of glycosylation across different cell lines (Fig. 3b). The variability in glycosylation patterns between cell lines underscores the challenges in glycoproteomics research. Gene Ontology (GO) analysis of proteins with high-frequency glycosylation (>75% of samples) revealed significant enrichment in cadherin binding pathways (Fig. 3c). Notably, proteins such as EGFR, CLINT1, ESYT2, ITGB1, EIF4G2, PLP4, TNND1, and PICALM, all heavily glycosylated, are involved in these pathways (Fig. 3d). For example, Epidermal Growth Factor Receptor (EGFR), a receptor tyrosine kinase and an important target for cancer therapies, was found to have 12 glycosylation sites (N1044, N1094, N128, N352, N361, N398, N413, N444, N526, N528, N592, N603, N623) (Fig. 3e). Among these, N361 and N352, located within the extracellular domain of EGFR, have been previously reported to be essential for maintaining EGF binding sites^31,47,48^. Additionally, N528 exhibits significant glycosylation, with 73 site-specific glycans and 44 different glycan types identified at this site. These modifications may play a critical role in the structure and function of EGFR, influencing its activity and interactions. Overall, our glycoproteomics into the glycosylation patterns of cancer cell lines helps to better understand critical drug targets and signaling pathways. This data presents an opportunity to explore how glycosylation and other PTMs, such as phosphorylation, interact to modulate protein function and cellular behavior, especially in receptor kinases like EGFR. Further targeted analyses could enhance our understanding of these interactions.

**Figure 3.**
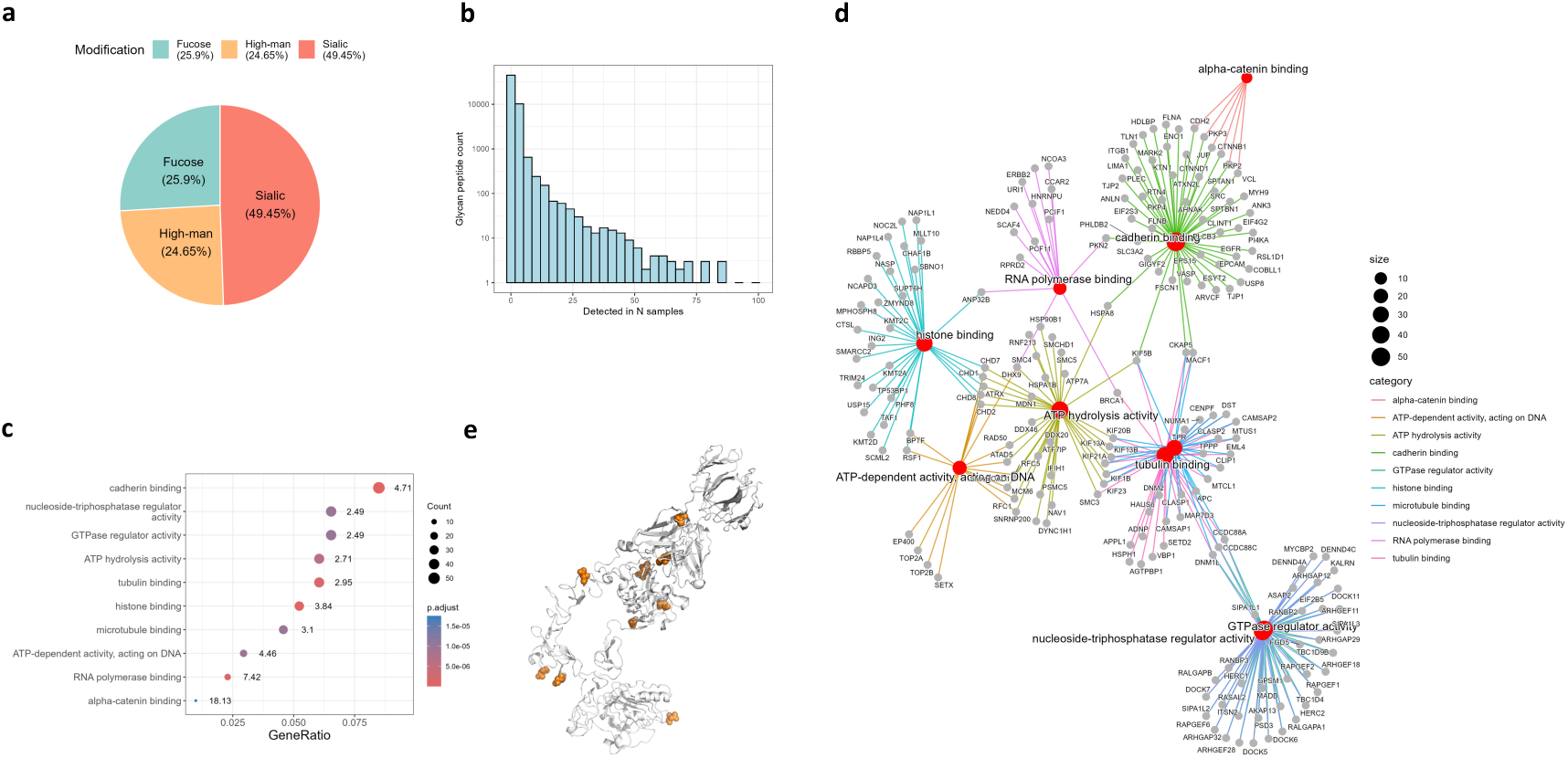
Glycoproteome Landscape of Cell Lines (a) Distribution of glycosylation types: Site-specific glycans were classified into sialic acid, fucose, and high-mannose types. A pie chart displays the percentage distribution of these glycan types across the cell lines. (b) Distribution of site-specific glycans: The Y-axis shows the counts of site-specific glycans identified by MS in the cell lines, while the X-axis represents the number of samples in which each glycopeptide was detected. (c) Enriched GO terms of heavily glycosylated proteins: Heavily glycosylated proteins were selected based on their occurrence in multiple cell lines. The p-value reflects the significance of the biological functions associated with these glycosylated proteins. (d) Glycosylated proteins involved in enriched functions: Highlights the specific proteins contributing to the enriched biological functions identified in panel (c). (e) Detected glycosylation sites on epidermal growth factor receptor (EGFR): Shows the glycosylation sites on EGFR detected across the different cell lines.

In conclusion, the glycosylation profiling of cancer cell lines provides a comprehensive view of glycoprotein status, offering a powerful tool for understanding the molecular behaviors of cancer cells and informing therapeutic strategies.

### Comparison of MS and RPPA proteomics data

The comparison between Mass Spectrometry (MS) and Reverse Phase Protein Array (RPPA) proteomics data reveals key insights into their performance and correlation. The Spearman correlation coefficients (Fig. 4a) show high intra-group consistency for both technologies, with values ranging from 0.85 to 0.95, indicating strong reproducibility within the same experimental conditions. Inter-group correlations are notably weaker, especially in phosphorylation datasets, where MS phosphoproteome inter-group correlations are 0.40, and RPPA phosphorylation sites are −0.007, showing greater variability across different cell lines. MS provides a detailed and comprehensive analysis of proteins and post-translational modifications (PTMs), including phosphorylation. However, it can exhibit higher variability between experimental groups and occasionally have missing data points. Conversely, RPPA is more standardized and reproducible, especially for high-throughput applications In this study, we used RPPA technology to analyze 231 proteins. Out of these, 212 proteins (91.8%) were successfully quantified by MS, but 19 proteins (8.2%) were missed by MS (Fig. 4c). Due to missing values in the MS data, only 146 proteins (∼63.2%) were quantified by MS in more than half of the samples, while 66 proteins (28.6%) were quantified in fewer than half. Functional analysis of these proteins revealed significant involvement in kinase and ubiquitase activity (Fig. 4d). When comparing the quantitative data from MS and RPPA, 90% of the proteins showed a positive correlation between the two methods, with a median correlation coefficient of 0.6 (Fig. 4e), indicating strong agreement between the technologies. Additionally, a comparison of protein fold changes between myeloid and tumor epithelial cells demonstrated a stronger correlation, with a Spearman coefficient of 0.79 (Fig. 4f), than the correlation between protein expression levels.

**Figure 4.**
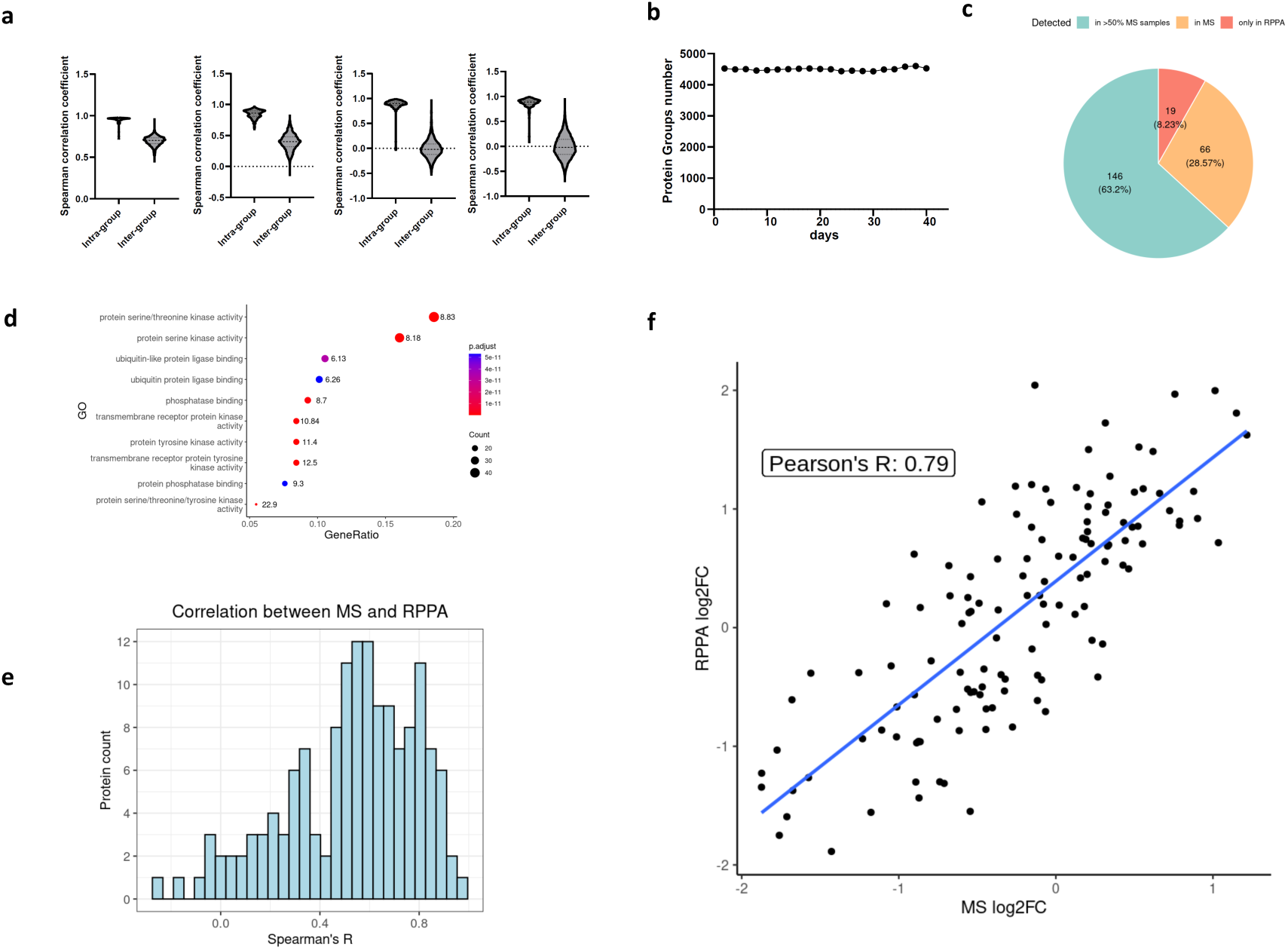
Comparison between RPPA and MS Whole Proteomics Data. (a) Spearman correlation coefficients: Comparison of intra- and inter-group correlations for MS whole proteome, MS phosphoproteome, RPPA whole proteins, and RPPA phosphorylation sites, shown from left to right. (b) MS instrument monitoring: Daily tracking of MS performance using whole-cell lysate (Hela cells) tryptic digest standards. (c) Protein detection comparison: Analysis of 231 proteins in the RPPA panel. Cyan indicates proteins detected by MS in more than 50% of samples, orange for proteins detected in fewer than 50%, and dark orange for proteins not detected by MS. (d) GO term enrichment: Functional analysis of proteins detected in RPPA proteomics, highlighting major biological processes associated with the RPPA panel. (e) Protein quantitation correlation: Spearman’s R-values show the correlation of protein levels between MS and RPPA data. The X-axis represents R-values, and the Y-axis shows the number of proteins within each range. (f) Fold change correlation: Comparison of protein expression fold changes across different cell lines as measured by MS and RPPA, based on datasets from myeloid and tumor epithelial cells.

In summary, MS and RPPA provide complementary insights into proteomics data. MS offers broad coverage of proteins and PTMs, though with occasional missing data points, while RPPA delivers more focused, consistent data based on antibody availability. Together, these methods validate and complement each other, confirming their reliability in quantitative proteomic analysis.

## Discussion

Previous studies have extensively profiled the whole proteome landscape of pan-cancer cell lines, offering valuable insights for biological research. However, these studies have largely overlooked the role of post-translational modifications (PTMs) such as phosphorylation and glycosylation, which are critical for processes like signal transduction, tumor progression, and drug responses. Our study addresses this gap by presenting comprehensive datasets focused on these key PTMs, creating a pan-cancer PTM landscape. With our phosphoproteomics data, we can identify activated signaling pathways and kinases across different cell lines, offering foundational data for selecting suitable models for experimental validation and clinical pharmacology. Additionally, our glycoproteomics data reveals glycosylation sites and glycan types on key signaling proteins, particularly on transmembrane receptors, enhancing our understanding of cellular regulation and protein functionality.

We used IMAC enrichment to capture both phosphorylated and glycosylated peptides, as IMAC is effective in capturing peptides containing phosphorous and sialic acid groups. The specific enrichment conditions favored phosphorylated peptides in this study. While the glycopeptides are less abundant than phosphopeptides, they serve as a valuable supplementary database without requiring extensive enrichment efforts or additional MS instrument time.

To facilitate quantifying site-specific glycans, we developed a novel method to quantify glycosylated protein expression based on spectral counting, the code for which is available in Supplemental Material 6. This code allows users to extract quantitative information from Byonic™ search results, bypassing the limitations of traditional glycopeptide identification tools that lack direct quantitation functionality. Our method simplifies the quantitation of relative abundances of site-specific glycans across different cancer cell lines.

We also compared MS with RPPA proteomics and found a strong correlation in the observed changes in protein expression between the two platforms. This consistency suggests that conclusions regarding the upregulation or downregulation of proteins across different sample groups remain reliable regardless of the technology used. MS offers broader coverage, detecting a wide range of proteins and PTMs, but can miss data for some proteins. RPPA, while more targeted, sensitive, and reproducible, is dependent on the availability of specific antibodies for the proteins and PTMs of interest, preventing it from being used in global proteome study.

In conclusion, our study systematically constructed the proteome, phosphoproteome, and glycoproteome landscape of human cancer cell lines, revealing consistent protein expression patterns across cancer types, identifying tissue-specific biomarkers, highlighting active signaling pathways, and detailing important protein modifications. These datasets are invaluable for studying cancer mechanisms, identifying biomarkers, and discovering potential therapeutic targets. Additionally, our comparison of MS and RPPA technologies provides guidance for choosing the most appropriate method in cancer proteomics research, depending on the experimental focus.

## Technical Validation

To ensure data reliability, we implemented stringent validation steps across the proteomics workflow:

1. High Intra-Group Correlation: The Spearman correlation coefficients calculated within the same cell lines show strong consistency for both MS and RPPA, with values ranging from 0.85 to 0.95. This indicates robust reproducibility under identical experimental conditions. In contrast, the correlations between different groups were low (Fig. 4a).
2. Mass Spectrometry (MS) Calibration: MS performance was validated daily using a HeLa cell lysate digest as a reference standard (Fig. 4b), ensuring consistent signal intensity and retention time stability throughout the study.
3. Correlation Validation: Protein fold changes between MS and RPPA showed strong agreement, with a Spearman correlation of 0.79 for protein perturbation (Fig. 4f), validating the accuracy across different technology platforms.

These validation steps confirm the robustness of our datasets, ensuring their suitability for future research in cancer biology.

## Authors and Affiliations

1. Chemistry Department, Tsinghua University, 30 Shuangqing Road, Haidian District, Beijing 100084 China Yiying Zhu, Wenhao Shi
2. Bioscience and Biomedical Engineering Thrust, Systems Hub, The Hong Kong University of Science and Technology (Guangzhou), Guangzhou, 511453 China. Division of Emerging Interdisciplinary Areas, Center for Aging Science, The Hong Kong University of Science and Technology, Clear Water Bay, Hong Kong, China. Tang Shaojun, He Tianlong, Annan Qian, Yuqiao Liu
3. Cosmos Wisdom Biotech Co. Ltd, Building 10th, No. 617 Jiner Road, Hangzhou, 311215, China. Nan Wang [This author was a previous employee of Mills Institute for Personalized Cancer Care, Fynn Biotechnologies, Jinan, China]

## Author contributions

Study conception and supervision: Y.Z., N.W., S.T., Investigation and acquisition of MS and RPPA proteomics data: W.S., N.W. Data analysis, integration, and interpretation: T.H., W.S., A.Q., Y.L., Y.Z. Supervision of the bioinformatics analysis: S.T. Writing: Y.Z., N.W., W.S., T.H. Reviewing and editing: Y.Z., N.W.

## Acknowledgment

We thank PRECEDO Biotechnologies (Hefei, China) for providing valuable cell lines for this work. We also thank other team members at Fynn Biotechnologies Co., Ltd (Shangdong, China) for conducting the RPPA.

## References

1 Nusinow, D. P. et al. Quantitative Proteomics of the Cancer Cell Line Encyclopedia. Cell 180, 387–402 e316 (2020). 10.1016/j.cell.2019.12.023

2 Guo, T. et al. Quantitative Proteome Landscape of the NCI-60 Cancer Cell Lines. iScience 21, 664–680 (2019). 10.1016/j.isci.2019.10.059

3 Gholami, A. M. et al. Global proteome analysis of the NCI-60 cell line panel. Cell Rep 4, 609–620 (2013). 10.1016/j.celrep.2013.07.018

4 Frejno, M. et al. Proteome activity landscapes of tumor cell lines determine drug responses. Nat Commun 11, 3639 (2020). 10.1038/s41467-020-17336-9

5 Coarfa, C. et al. Reverse-Phase Protein Array: Technology, Application, Data Processing, and Integration. J Biomol Tech 32, 15–29 (2021). 10.7171/jbt.21-3202-001

6 Ding, Z., Wang, N., Ji, N. & Chen, Z. S. Proteomics technologies for cancer liquid biopsies. Mol Cancer 21, 53 (2022). 10.1186/s12943-022-01526-8

7 Wang, N. et al. Parallel Analyses by Mass Spectrometry (MS) and Reverse Phase Protein Array (RPPA) Reveal Complementary Proteomic Profiles in Triple-Negative Breast Cancer (TNBC) Patient Tissues and Cell Cultures. 2024.2005.2030.596640 (2024). 10.1101/2024.05.30.596640 %J bioRxiv

8 Lei, Z. N. et al. ABCB1-dependent collateral sensitivity of multidrug-resistant colorectal cancer cells to the survivin inhibitor MX106-4C. Drug Resist Updat 73, 101065 (2024). 10.1016/j.drup.2024.101065

9 Shankavaram, U. T. et al. Transcript and protein expression profiles of the NCI-60 cancer cell panel: an integromic microarray study. Mol Cancer Ther 6, 820–832 (2007). 10.1158/1535-7163.MCT-06-0650

10 Park, E. S. et al. Integrative analysis of proteomic signatures, mutations, and drug responsiveness in the NCI 60 cancer cell line set. Mol Cancer Ther 9, 257–267 (2010). 10.1158/1535-7163.MCT-09-0743

11 Tian, Q. et al. Integrated genomic and proteomic analyses of gene expression in Mammalian cells. Mol Cell Proteomics 3, 960–969 (2004). 10.1074/mcp.M400055-MCP200

12 Federici, G. et al. Systems analysis of the NCI-60 cancer cell lines by alignment of protein pathway activation modules with “-OMIC” data fields and therapeutic response signatures. Mol Cancer Res 11, 676–685 (2013). 10.1158/1541-7786.MCR-12-0690

13 Li, J. et al. Characterization of Human Cancer Cell Lines by Reverse-phase Protein Arrays. Cancer Cell 31, 225–239 (2017). 10.1016/j.ccell.2017.01.005

14 Ghandi, M. et al. Next-generation characterization of the Cancer Cell Line Encyclopedia. Nature 569, 503–508 (2019). 10.1038/s41586-019-1186-3

15 Vizcaino, J. A. et al. The PRoteomics IDEntifications (PRIDE) database and associated tools: status in 2013. Nucleic Acids Res 41, D1063–1069 (2013). 10.1093/nar/gks1262

16 Li, J. et al. A protein expression atlas on tissue samples and cell lines from cancer patients provides insights into tumor heterogeneity and dependencies. Nat Cancer 5, 1579–1595 (2024). 10.1038/s43018-024-00817-x

17 Meissner, F., Geddes-McAlister, J., Mann, M. & Bantscheff, M. The emerging role of mass spectrometry-based proteomics in drug discovery. Nat Rev Drug Discov 21, 637–654 (2022). 10.1038/s41573-022-00409-3

18 Nan Wang, Y. Z., Lianshui Wang, Wenshuang Dai, Taobo Hu, Zhentao Song, Xia Li, Qi Zhang, Jianfei Ma, Qianghua Xia, Jin Li, Yiqiang Liu, Mengping Long, Zhiyong Ding. (bioRxiv, 2024).

19 Tibes, R. et al. Reverse phase protein array: validation of a novel proteomic technology and utility for analysis of primary leukemia specimens and hematopoietic stem cells. Mol Cancer Ther 5, 2512–2521 (2006). 10.1158/1535-7163.MCT-06-0334

20 Wang, N. et al. A reverse phase protein array based phospho-antibody characterization approach and its applicability for clinical derived tissue specimens. Sci Rep 12, 22373 (2022). 10.1038/s41598-022-26715-9

21 Abrey, L. E. et al. Report of an international workshop to standardize baseline evaluation and response criteria for primary CNS lymphoma. J Clin Oncol 23, 5034–5043 (2005). 10.1200/JCO.2005.13.524

22 Johansson, H. J. et al. Breast cancer quantitative proteome and proteogenomic landscape. Nat Commun 10, 1600 (2019). 10.1038/s41467-019-09018-y

23 Savage, S. R. et al. Pan-cancer proteogenomics expands the landscape of therapeutic targets. Cell 187, 4389–4407 e4315 (2024). 10.1016/j.cell.2024.05.039

24 Roskoski, R., Jr. Properties of FDA-approved small molecule protein kinase inhibitors: A 2024 update. Pharmacol Res 200, 107059 (2024). 10.1016/j.phrs.2024.107059

25 Roskoski, R., Jr. Cost in the United States of FDA-approved small molecule protein kinase inhibitors used in the treatment of neoplastic and non-neoplastic diseases. Pharmacol Res 199, 107036 (2024). 10.1016/j.phrs.2023.107036

26 Wang, Y. et al. FDA-approved small molecule kinase inhibitors for cancer treatment (2001-2015): Medical indication, structural optimization, and binding mode Part I. Bioorg Med Chem 111, 117870 (2024). 10.1016/j.bmc.2024.117870

27 Pinho, S. S. & Reis, C. A. Glycosylation in cancer: mechanisms and clinical implications. Nat Rev Cancer 15, 540–555 (2015). 10.1038/nrc3982

28 Cohen, M. H. et al. Approval summary for imatinib mesylate capsules in the treatment of chronic myelogenous leukemia. Clin Cancer Res 8, 935–942 (2002).

29 Rikova, K. et al. Global survey of phosphotyrosine signaling identifies oncogenic kinases in lung cancer. Cell 131, 1190–1203 (2007). 10.1016/j.cell.2007.11.025

30 He, K. et al. Decoding the glycoproteome: a new frontier for biomarker discovery in cancer. J Hematol Oncol 17, 12 (2024). 10.1186/s13045-024-01532-x

31 Lih, T. M., Cho, K. C., Schnaubelt, M., Hu, Y. & Zhang, H. Integrated glycoproteomic characterization of clear cell renal cell carcinoma. Cell Rep 42, 112409 (2023). 10.1016/j.celrep.2023.112409

32 Hu, Y., Shah, P., Clark, D. J., Ao, M. & Zhang, H. Reanalysis of Global Proteomic and Phosphoproteomic Data Identified a Large Number of Glycopeptides. Anal Chem 90, 8065–8071 (2018). 10.1021/acs.analchem.8b01137

33 Cho, K. C., Chen, L., Hu, Y., Schnaubelt, M. & Zhang, H. Developing Workflow for Simultaneous Analyses of Phosphopeptides and Glycopeptides. ACS Chem Biol 14, 58–66 (2019). 10.1021/acschembio.8b00902

34 Meng, Q., Xia, C., Fang, J., Rojanasakul, Y. & Jiang, B. H. Role of PI3K and AKT specific isoforms in ovarian cancer cell migration, invasion and proliferation through the p70S6K1 pathway. Cell Signal 18, 2262–2271 (2006). 10.1016/j.cellsig.2006.05.019

35 Khabele, D. et al. Preferential effect of akt2-dependent signaling on the cellular viability of ovarian cancer cells in response to EGF. J Cancer 5, 670–678 (2014). 10.7150/jca.9688

36 Huang, Q. et al. Akt2 kinase suppresses glyceraldehyde-3-phosphate dehydrogenase (GAPDH)-mediated apoptosis in ovarian cancer cells via phosphorylating GAPDH at threonine 237 and decreasing its nuclear translocation. J Biol Chem 286, 42211–42220 (2011). 10.1074/jbc.M111.296905

37 Noske, A. et al. Specific inhibition of AKT2 by RNA interference results in reduction of ovarian cancer cell proliferation: increased expression of AKT in advanced ovarian cancer. Cancer Lett 246, 190–200 (2007). 10.1016/j.canlet.2006.02.018

38 Yuan, Z. Q. et al. Frequent activation of AKT2 and induction of apoptosis by inhibition of phosphoinositide-3-OH kinase/Akt pathway in human ovarian cancer. Oncogene 19, 2324–2330 (2000). 10.1038/sj.onc.1203598

39 Tanaka, Y. et al. Abemaciclib, a CDK4/6 inhibitor, exerts preclinical activity against aggressive germinal center-derived B-cell lymphomas. Cancer Sci 111, 749–759 (2020). 10.1111/cas.14286

40 Pikman, Y. et al. Synergistic Drug Combinations with a CDK4/6 Inhibitor in T-cell Acute Lymphoblastic Leukemia. Clin Cancer Res 23, 1012–1024 (2017). 10.1158/1078-0432.CCR-15-2869

41 Bride, K. L. et al. Rational drug combinations with CDK4/6 inhibitors in acute lymphoblastic leukemia. Haematologica 107, 1746–1757 (2022). 10.3324/haematol.2021.279410

42 Borgo, C., D’Amore, C., Sarno, S., Salvi, M. & Ruzzene, M. Protein kinase CK2: a potential therapeutic target for diverse human diseases. Signal Transduct Target Ther 6, 183 (2021). 10.1038/s41392-021-00567-7

43 Prins, R. C. et al. CX-4945, a selective inhibitor of casein kinase-2 (CK2), exhibits anti-tumor activity in hematologic malignancies including enhanced activity in chronic lymphocytic leukemia when combined with fludarabine and inhibitors of the B-cell receptor pathway. Leukemia 27, 2094–2096 (2013). 10.1038/leu.2013.228

44 Zhang, L. et al. Comprehensive landscape of gastric cancer-targeted therapy and identification of CSNK2A1 as a potential target. Heliyon 10, e36205 (2024). 10.1016/j.heliyon.2024.e36205

45 Schjoldager, K. T., Narimatsu, Y., Joshi, H. J. & Clausen, H. Global view of human protein glycosylation pathways and functions. Nat Rev Mol Cell Biol 21, 729–749 (2020). 10.1038/s41580-020-00294-x

46 Riley, N. M., Bertozzi, C. R. & Pitteri, S. J. A Pragmatic Guide to Enrichment Strategies for Mass Spectrometry-Based Glycoproteomics. Mol Cell Proteomics 20, 100029 (2021). 10.1074/mcp.R120.002277

47 Lam, D., Arroyo, B., Liberchuk, A. N. & Wolfe, A. L. Effects of N361 Glycosylation on Epidermal Growth Factor Receptor Biological Function. bioRxiv (2024). 10.1101/2024.07.12.603279

48 Azimzadeh Irani, M., Kannan, S. & Verma, C. Role of N-glycosylation in EGFR ectodomain ligand binding. Proteins 85, 1529–1549 (2017). 10.1002/prot.25314

